# Attentional cueing effects are reversed during locomotion

**DOI:** 10.1101/2025.01.07.631651

**Authors:** Zakaria Djebbara, Dylan Chau, Aleksandrs Koselevs, Yiru Chen, Lars Brorson Fich, Klaus Gramann

**Affiliations:** Department of Architecture, Design, Media and Technology, Aalborg University, Aalborg, Denmark; Biological Psychology and Neuroergonomics, Technische Universität Berlin, Berlin, Germany

**Author notes:** **Conflict of interest.** The authors declare no conflict of interest. **Source data.** All data is available on: https://www.osf.io/92vzx/.

**Keywords:** Mobile Neuroimaging, Mobile Brain/Body Imaging, Architectural Cognition, Exogenous Attention, Posner Paradigm, Transitions, Naturalistic Paradigms

## Abstract

Everyday human cognition and behaviour evolved in dynamic and ever-changing environments, but static paradigms still dominate experimental research despite concerns about generalisability of the results. In the case of attention, traditional stationary studies show that pre-orienting attention with spatial cues leads to faster, more accurate responses. However, how movement and environmental features shape such attentional processes in everyday behaviour remains unknown. Here we show that active movement through curved corridors reverses the typical spatial attention effect, with faster response times and higher accuracy for stimuli incongruent to implicit spatial cues provided by the movement direction, contradicting previous findings from static settings. We found that early (N1) and late (P3) attention-related electrophysiological responses were modulated by environmental features and motor demands. The posterior N1-component, reflecting visuo-spatial attention, showed decreasing amplitudes as turning angles and motor-control demands increased for congruent stimuli appearing on the side of the turning direction. Similarly, the P3-complex varied with motor and visual processing demands for congruent stimuli, showing decreased amplitudes as motor-control demands increased. We propose that congruent stimuli, displayed against a dynamically changing visual context, increase pulvino-cortical processing load and slowing early visual processing that affect behavioural responses. Incongruent stimuli, however, are displayed against a predictable context allowing faster target processing. These findings challenge attentional mechanisms’ assumed consistency across static and dynamic settings, revealing instead their dependence on behavioural and environmental context. We advocate for naturalistic paradigms, arguing that moving beyond static experiments could reshape core views on cognition and behaviour.

## 1. Introduction

In everyday life, we are naturally exposed to a multitude of ongoing stimuli. Our attentional system selectively focuses on selected streams of information whilst suppressing others ^1,2^. While the neural dynamics of our attentional system have been studied using static experimental setups with participants seated, or lying in controlled environments, very little has been done to understand this system in natural, active circumstances. Laboratory studies are criticised because they avoid real-world settings where attentional shifts, decision-making, and embodied interactions are shaped by an ever-changing world. This has been met by a call for naturalistic paradigms in cognitive neuroscience as neural responses are to a large extent interrelated with behaviour, body, and environment ^3–7^. This is further supported theoretically from the increasing interest in understanding cognitive skills as embedded, extended, embodied and enacted ^6,8–12^, typically referred to as 4E Cognition. Such naturalistic settings can reveal how the brain truly functions in dynamic, everyday situations ^13,14^. Should naturalistic experiments reveal divergent neural and behavioural patterns compared with those observed in the laboratory, it becomes imperative to critically evaluate the extent to which existing lab-based findings generalise to real-world contexts.

The present study adapted a classic attentional cueing paradigm to a more natural setting by using corners in the built environment, enabling us to examine how turning directions exogenously cue attention. We used a *Mobile Brain/Body Imaging* approach (MoBI; ^5,15,16^), which allows for studying the synchronised behavioural and neural dynamics underlying human cognition in this kind of naturalistic settings. Specifically, we adapted a Posner cueing paradigm ^17^ where human participants physically walked through corridors with varying angles in Virtual Reality (VR), where the turning direction acts as the implicit spatial cue as small spheres appeared systematically to the same or the opposite side as the turning direction. Cueing paradigms result in faster and more accurate responses when the target stimulus is displayed on the same side as the cue, i.e., on congruent trials ^17–19^. The response advantage is taken as evidence that attention can be pre-oriented through covert attention strategies ^2,20^. However, these effects have so far only been observed under highly controlled experimental conditions, where participants are required to remain seated or lying down and refrain from moving their eyes or head toward the stimuli.^21^. This contrasts sharply with everyday interactions where people constantly move their eyes, head, and body, actively shifting their attention ^9,22–25^.

In this study, we modified the traditional cueing paradigm to increase its ecological validity. In typical stationary protocols, cues are presented as static indicators—like an arrow or a brief change in color or shape—that precede the target stimulus and often predict its location. By contrast, continuous cues, such as optic flow, provide an ongoing signal that encourages sustained and fluid shifts of attention, reflecting how attention naturally operates in dynamic, real-world contexts. In addition, to examine how active movement through the space shapes attentional processes, the cues were made non-informative regarding the target location, i.e., they appeared at chance level on the same or opposite side of the turning direction. This way, we introduced ecological visual and motor signals to investigate their impact on human brain responses underlying spatial attention during active locomotion. Recent attention studies suggest that the thalamus, particularly the pulvinar, serves as an integrative hub between visual and motor signals, establishing cortico-pulvino-cortical loops that connect various cortical areas, including the striate and extrastriate cortices ^26–34^. By integrating top-down predictions and bottom-up prediction-errors ^35^, the pulvinar plays a key role in shaping saliency maps that aid visual attention and the coordination of visually guided movements ^36,37^. Importantly, neurons in the pulvinar exhibit positive correlations with optic flow, indicating that the pulvinar may be sensitive to variations in sensory velocity (optic flow), which play a significant role in motor control ^38–41^. This integrative view of the pulvinar portrays it as a critical mediator in the flow of information between sensory inputs and motor outputs, and as critical to shaping perceptual and cognitive functions ^42–44^.

Our daily movement through built environments involves varying demands from both the task and surrounding conditions, as reflected by gaze strategies that prioritise task-relevant information ^45–47^. For example, head-and eye-tracking studies show that when turning a corner, people consistently shift their gaze in the same direction as the turn ^45–51^. Sometimes referred to as gaze-cueing, this mechanism reliably directs observers’ attention ^21^. Compelling evidence suggests that attention is guided by gaze actions that minimize uncertainty—or Bayesian surprise—in the visual field ^52–55^. When turning to the right, for example, individuals tend to orient their gaze in the same direction as this provides novel information ^51^. Thus, turning corners naturally operationalizes an attentional cueing paradigm (Figure 1). Studies on human gait further indicate that people make turning decisions that conserve energy ^56–58^, even if such decisions may occasionally lead to less favourable outcomes ^59^. Because sharper turns demand more energy ^60,61^, it follows that attention may be influenced by the physical effort required by the built environment. In this way, turning angles become a valuable factor for exploring how perceptual, attentional, and motor processes are shaped by our surroundings— underscoring the importance of the built environment in cognition and behaviour ^62–65^.

**Figure 1.**
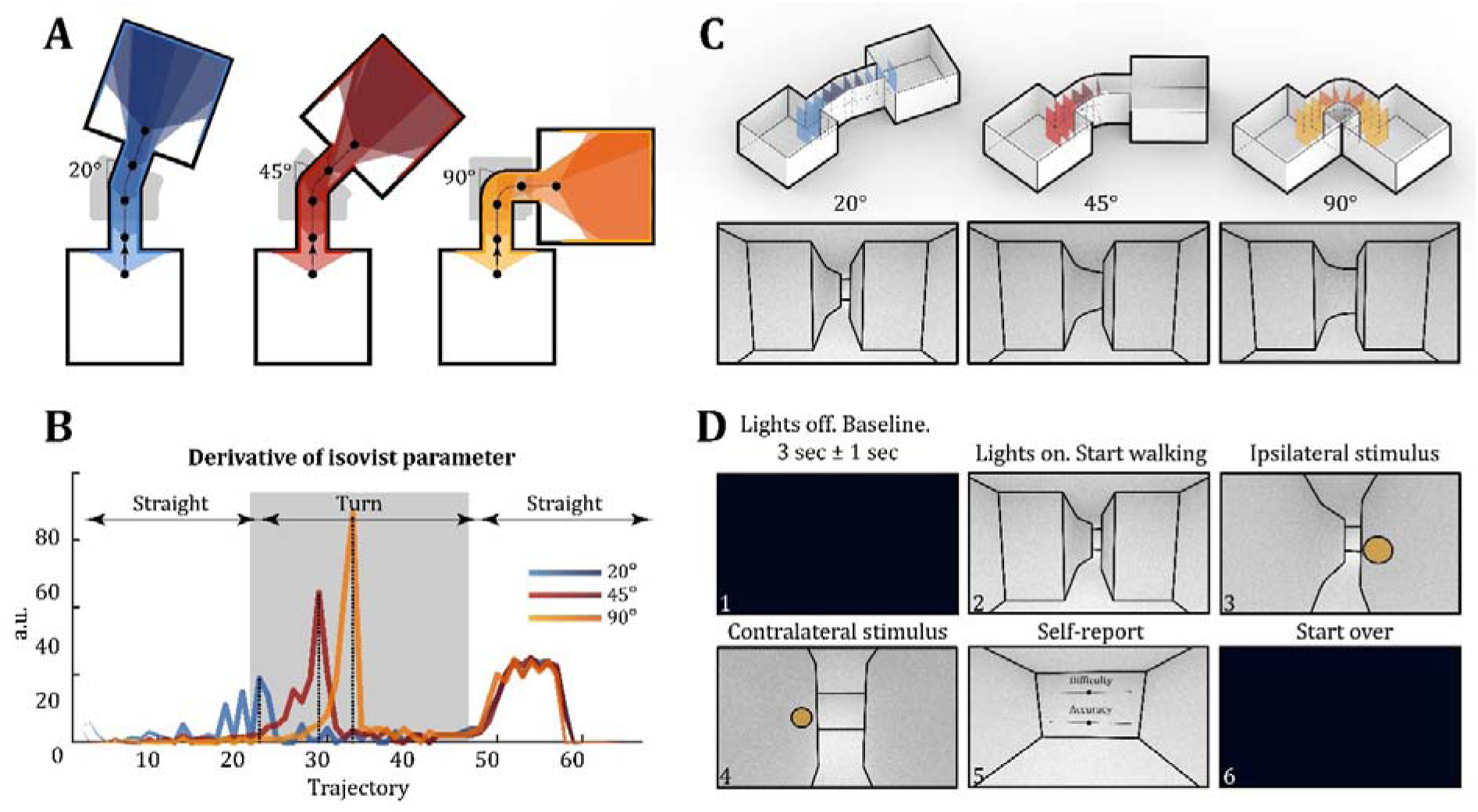
**A.** Three layouts of the virtual environment used in the study and an Isovist agent (see Methods) passing through the corner. The coloured walls illustrate the visual contact obtained by the agent (field of view: 120 degrees). The grey area coloured at the centre refers to the *Turning* phase, which captures the actual turning of the corner. **B.** By quantifying the visual contact, the velocity of novel visual information can be computed. These graphs illustrate the respective sensory velocity of each corner, showcasing that turning a 90° corner results in receiving lots of new visual information in relatively short time. The background of the graph illustrates where the *Turning* and *Straight* phases are located relative to the sensory velocity. **C.** Three 3D-layouts (without roof) displaying the three virtual environments for right-turn condition, as an example. The coloured walls are invisible to the participants and acts as triggers for the stimuli. The *Turning* phase is coloured in slightly darker colours in each respective condition. **D.** The experimental paradigm starts in the dark, providing us with a baseline for 3 ± 1 second. When the lights turn on, participants are met with one of the six conditions, i.e., a 20°, 45° or 90° corner, turning either to the right or left. They are instructed to begin walking and respond (by pulling the trigger of the Virtual Reality controller) according to the direction in which the stimulus appears. In the final space, they briefly report on the perceived difficulty and accuracy before the lights turn off again.

Essentially, this study is a naturalised Posner paradigm with human participants walking through VR-environments with corners of varying degrees while their neuronal activity is continuously measured. We analysed the event-related potentials (ERPs) with onset of target stimuli as well as response to the same along the paths. The turning directions functioned as attentional cues while the angles of the corners manipulated the difficulty of the motor task. We hypothesised (1) to find faster response times and higher accuracy in congruent trials, i.e. stimuli on the same side as the turning direction. Similarly, we expected (2) to find significantly slower response times and lower accuracy during the physical act of the turn due to greater distribution of neural resources ^66^. For early event-related potentials of the electroencephalogram (EEG), we expected (3) to observe more pronounced N1 amplitudes to stimuli displayed in the visual field congruent with the turning direction due to the orientation of attention ^67–69^. We also expected to find attenuated P3 amplitudes for sharper turns due to the distribution of cognitive resources in more physically demanding settings ^70,71^.

## 2. Results

Twenty participants (9 females), between 23 and 54 years of age (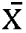= 30.1, σ= 7.18), performed the task in VR. While walking through virtual corridors, a total of 10 small spheres appeared on either the same or opposite side of the turning direction, i.e., in a congruent or incongruent position, respectively. The onset of spheres was elicited by invisible colliders in VR in space making them location-dependent, rather than time-dependent. Participants performed 240 trials, where each consisted of 10 pseudorandomised stimuli (5 stimuli on each side). A 90.38% of all trials were correctly answered, leaving 9.62% incorrectly answered. After preprocessing the data, a total of 37.882 trials entered the analyses.

### a. Turning corners over 30° affect the cognitive load

To answer whether the degree and the direction of turning biased the reaction times (RTs), we performed both a linear and generalised mixed-effects regression analysis, including the factors *Turning Angle* (20°,45°,90°), *Congruence* (ipsi-, contralateral stimuli relative to turning direction), and the *Stimulus Position* (1 through 10). This generated poorly fitted models that explained 38.2% and 1.2% of the variance, respectively, as they did not account for differences in RTs during straight aspects of the passage and the physical turn (Figure 2A; note the bump between Stimuli 4 and 7. See Figure 2B for separated conditions).

**Figure 2.**
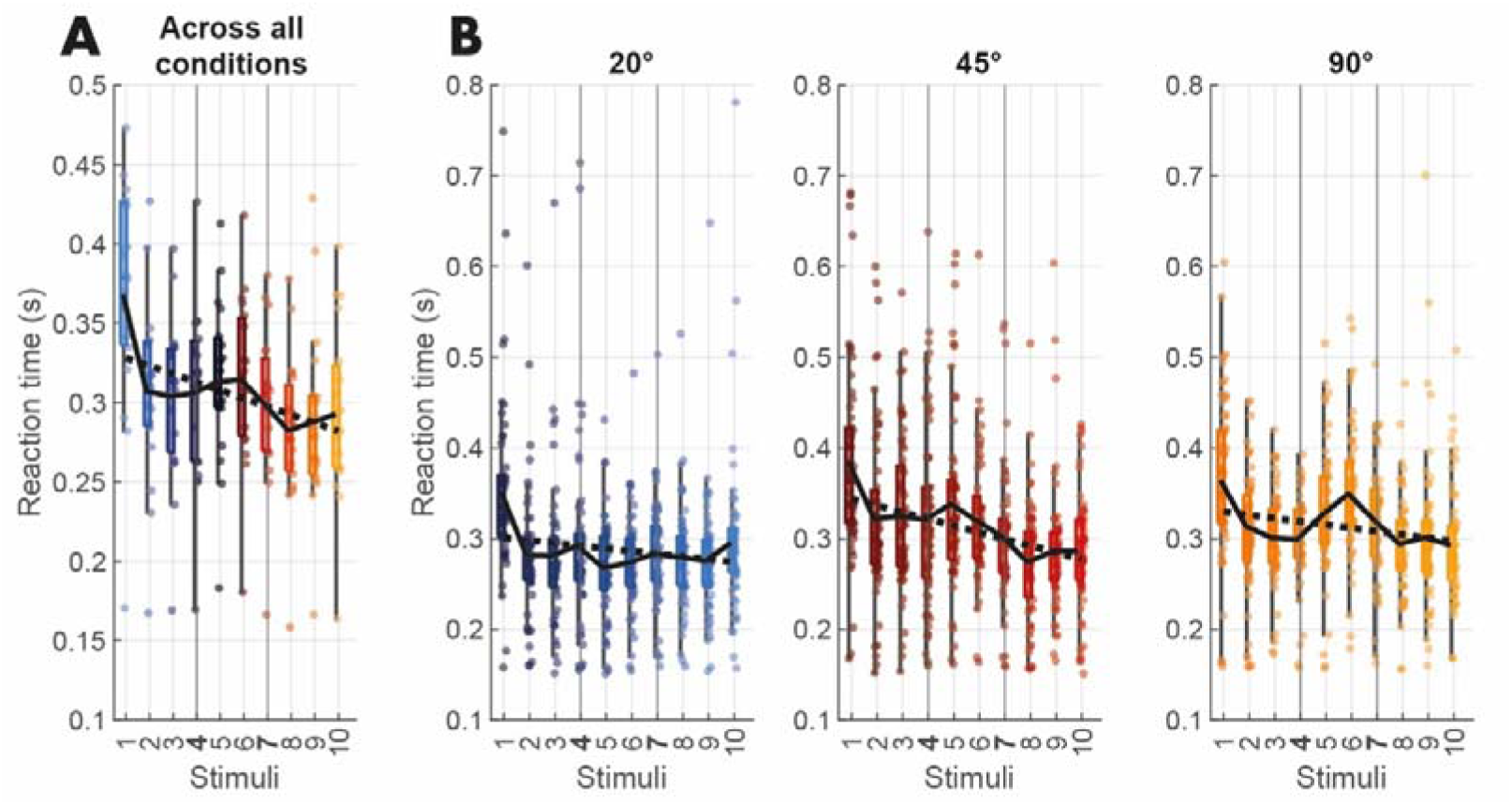
**A.** Recorded reaction times (depicted as dots; mean is depicted as solid line) of all *Turning Angles* plotted against the fitted mixed-effects model (depicted as the dashed line). Stimuli 4 and 7 represent the threshold between the *Straight* and *Turning* phases. Stimuli 1 through 3 and stimuli 8 through 10 are considered part of the *Straight* phase. Stimuli 4 through 7 are considered part of the *Turning* phase. **B.** These plots depict the same as A but split into their respective conditions. Notice the bump in the *Turning* phase particularly for 45° and 90° turns.

Accounting for the different phases by categorising and collapsing the *Stimulus Position* into *Phases*, which included *Turning* (stimuli 4 through 7) and *Straight* parts of the path (stimuli 1 through 3 and 8 through 10), we then recomputed the mixed-effects model where both models greatly improved (RTs: R^2^ = 89.5%; CRs: R^2^ = 88.7%), emphasising the impact of the turns on behaviour. The ANOVA of the mixed-effects model of RTs (Figure 3A and 3B) revealed significant main effects of the *Turning Angle* (F_2,221_ = 67.47; p < 0.001;η*_p_*^2^ = 0.2339) and the *Congruence* (F_1,221_ = 4.75; p = 0.0302;η*_p_*^2^ = 0.0210), but not for the *Phases* (F_1,221_ = 2.36; p = 0.1253;η*_p_*^2^ = 0.0106). However, we found significant interaction effects between both *Turning Angle* and *Congruence* (F2,221 = 7.09; p = 0.0083;η*_p_*^2^ = 0.0311) and *Turning Angle* and *Phases* (F2,221 = 15.57; p = 0.0001;η*_p_*^2^ = 0.0658). All significant factors required post-hoc analyses for better understanding.

**Figure 3.**
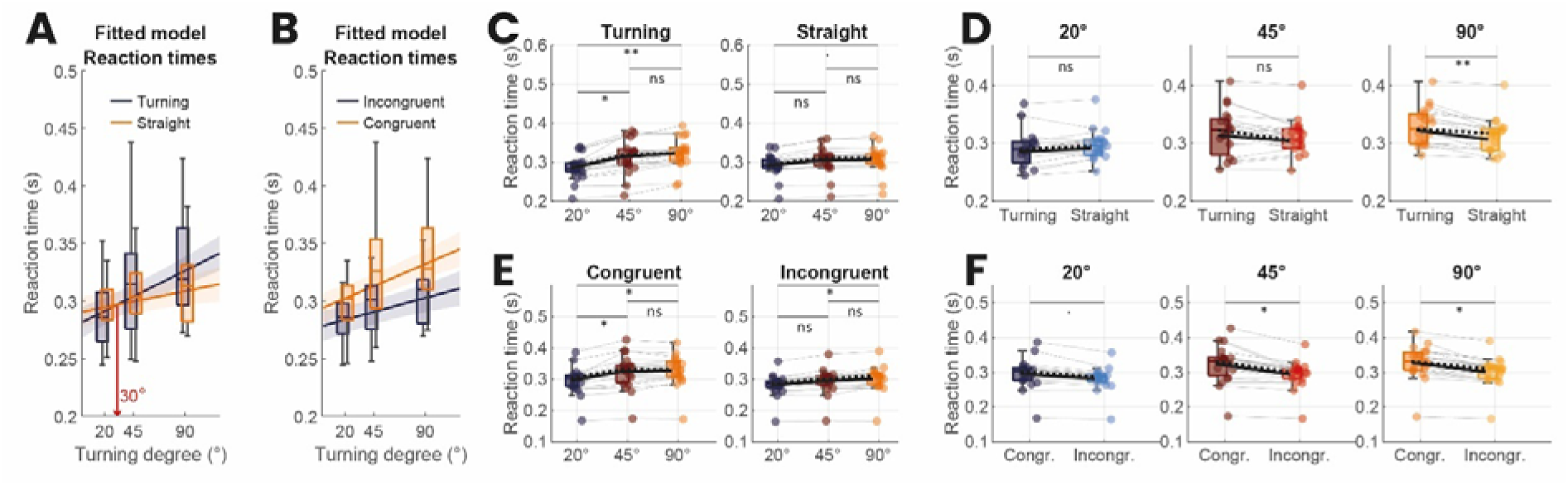
**A.** Plot of the fitted model in the case of *Phase*. Model estimates are plotted as solid lines, where the shaded areas indicate 95% confidence intervals. Boxplots refer to measured reaction times for each condition. We found that the *Straight* phase crosses the *Turning* phase at 30°, indicating this is where *Turning* generated longer RTs compared to *Straight* phases. **B.** Plot of the fitted model in the case of *Congruence*. **C.** Raincloud plots categorised by the *Phases*. We found increasing RTs for increasing degrees of turns, and we only found differences for *Turning*. **D.** Raincloud plots grouped by *Turning Angle*. We found increasing RTs for increasing difficulty. **E.** Here we contrasted the *Turning Angle* as grouped by *Congruence*. **F.** Finally, we contrasted the *Congruence* as grouped by *Turning Angle*. To our surprise, we found increasing RTs for *Congruent* stimuli. All contrasts here were computed using Wilcoxon Signed Rank test and corrected for multiple comparison using Benjamini-Hochberg.

Wilcoxon signed rank test (Figure 3C) revealed significantly faster RTs for 20° compared to 45° turns (p = 0.0369; d = 0.74), and even more so for 90° turns (p = 0.0039; d = 1.04). There were no significant differences between 45° and 90° (p = 0.4137; d = 0.21). We also could not identify significant differences for any *Straight Phases* for passages with different *Turning Angle*: 20° against 45° (p = 0.1197; d = 0.4), 20° against 90° (p = 0.0921; d = 0.47), and 45° against 90° (p = 0.7261; d = 0.07).

Comparing the *Phases* within each *Turning Angle* (Figure 3D), we only found significantly faster RTs for straight as compared to *Turning* phases for 90° turns (p = 0.0011; d = 0.33). There were no differences for 20° (p = 1175; d = 0.14) nor 45° (0.1815; d = 0.19).

For *Congruent* stimuli (Figure 3E), contrasts between 20° and 45° turns (p = 0.0496; d = 0.56) and between 20° and 90° (p = 0.0306; d = 0.67) reached significance, but not between 45° and 90° (p = 0.7481; d = 0.07). For *Incongruent* stimuli, significant differences were only identified between 20° and 90° turns (p = 0.0426; d = 0.50). No effects were observed between 20° and 45° (p = 0.1123; d = 0.32), and between 45° and 90° (p = 0.2933; d = 0.17). Comparing *Congruences* within *Turning Angle* revealed increasing effects on RTs with increasing turning degrees. Surprisingly, we also discovered that *Incongruent* stimuli resulted in faster RTs (Figure 3F). *Congruent*, compared to *Incongruent*, stimuli resulted in significantly slower RTs for both 45° (p = 0.0167; d = 0.61) and 90° (p = 0.0268; d =0.57), but not for 20° (p = 0.0961; d = 0.34).

These results suggest a threshold where the degree of the turn begins to affect reaction times more strongly, with our model identifying this threshold at 30°. Turns greater than 30° significantly slow down reaction times compared to straight paths, and, surprisingly, *Incongruent* stimuli generate increasingly faster reaction times than *Congruent* ones as turning angles increase.

### b. Enhanced performance for contralateral, not ipsilateral, stimuli

To assess the impact on the number of correct responses (CRs), we also fitted a mixed-effects model (Figure 4A and 4B). The ANOVA of the mixed-effects model revealed significant main effects for *Turning Angle* (F_2,221_ = 211.46, p < 0.0001, η*_p_*^2^ = 0.48), *Phases* (F_1,221_ = 39.02, p < 0.0001, η*_p_*^2^ = 0.15), and for *Congruence* (F_1,221_ = 12.93, p = 0.0003, η*_p_*^2^ = 0.05). There were also significant interaction effects between *Turning Angle* and *Congruence* (F_1,221_ = 91.53, p < 0.0001, η*_p_*^2^ = 0.29), between *Turning Angle* and *Phases* (F_1,221_ = 24.8, p < 0.0001, η*_p_*^2^ = 0.10), and between *Phases* and *Congruence* (F_1,221_ = 6.14, p = 0.0139, η*_p_*^2^ = 0.02).

**Figure 4.**
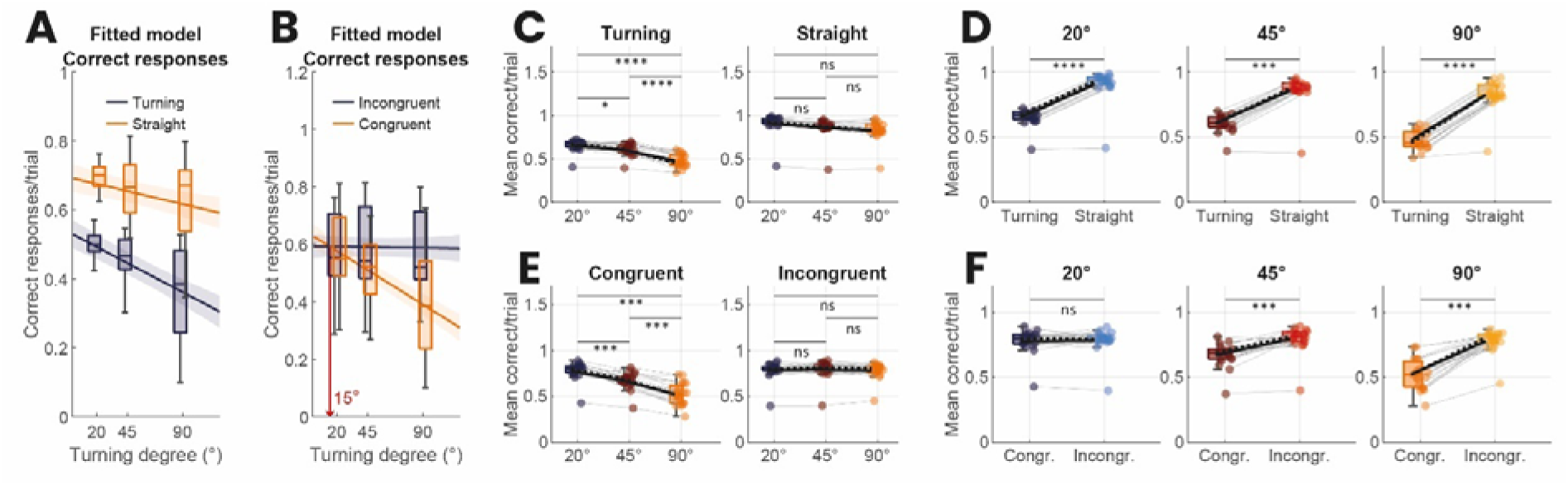
**A.** Plot of the fitted model in the case of *Phases*. Model estimates are plotted as solid lines, where the shaded areas indicate 95% confidence intervals. Boxplots refer to correct responses for each condition. **B.** Plot of the fitted model in the case of *Congruence*. We found that *Congruent* crosses *Incongruent* stimuli at 15°, suggesting that this is where *Congruent* stimuli generate more incorrect responses compared to *Incongruent* stimuli. **C.** Raincloud plots between the *Turning Angle* grouped by *Phases*. We found worsening performance for increasing degrees of turns, exclusively for the *Turning Phase*. **D.** Raincloud plots depict between the *Phases* grouped by *Turning Angle*. We found robustly worse performance during *Turning*. **E.** Here we contrasted the *Turning Angle* as grouped by *Congruence*. We found worsening performance for increasing *Turning Angle*, but exclusively for *Congruent* stimuli. **F.** Finally, we contrasted *Congruence* as grouped by *Turning Angle*, which revealed worsening performance for *Congruent* stimuli. All contrasts here were computed using Wilcoxon Signed Rank test and corrected for multiple comparison using Benjamini-Hochberg.

For *Turning*, post-hoc comparisons (Figure 4C) revealed significantly fewer CRs for 90° than 20° (p < 0.0001; d = 2.50) and 45° turns (p < 0.0001; d = 1.81), but also between 20° and 45° (p = 0.0431; d = 0.68). No significant effects emerged for the *Straight* phase: neither between 20° and 45° (p = 0.3402; d = 0.39), 20° and 90° (p = 0.1338; d = 0.67), nor between 45° and 90° (p = 0.3996; d = 0.27). However, we found that *Turning* resulted in consistently fewer CRs when compared with straight phases of the passages for each of the different *Turning Angles* (Figure 4D) - for 20° (p = 0.0001; d = 2.56), 45° (p = 0.0002; d = 2.56), and 90° (p = 0.0001; d = 3.51). Thus, CRs depended on the degree of the turn but also on the phase of the passage, where the physical turning caused significantly fewer CRs.

For *Congruency*, we found that CRs only differed for the *Congruent* stimuli (Figure 4E). We observed significantly more CRs for 20° compared to 45° (p = 0.0004; d = 1.12) and 90° (p = 0.0003; d = 2.27), but also for 45° and 90° (p = 0.0003; d = 1.30). No significant differences were observed for the *Incongruent* stimuli: neither between 20° and 45° (p = 0.5640; d = 0.11), 20° and 90° (p = 0.9359; d = 0.02), nor between 45° and 90° (p = 0.2106; d = 0.08). However, when comparing the *Congruence* within each *Turning Angle* (Figure 4F), we found significant differences for the 45° (p = 0.0006; d = 1.23), and 90° (p = 0.0003; d = 2.42), but not for 20° (p = 1; d = 0.04).

In sum, we found that participants scored increasingly worse for *Congruent* stimuli as the degree of turns increased, while the turning angles had no effect on *Incongruent* stimuli. This indicates that there is a threshold at which the degree of turning significantly impacts the CRs, with our model identifying this threshold at 15°. That is, the effect of the stimulus being *Congruent* or *Incongruent* becomes significant for turns exceeding 15°.

### c. N1 and P3 reflect behavioural performance

The behavioural results suggest that (1) the sharper the physical turn, the slower the response times for both *Congruent* and *Incongruent* stimuli, but more so for *Congruent* stimuli, and (2) the sharper the physical turn, the more incorrect responses are given, again most pronounced for *Congruent* stimuli. While we expected behavioural compromises for the sharp turns, we did not anticipate that this would be expressed more strongly for *Congruent* conditions, contradicting the traditional cueing results.

To understand the observed behaviour, we further analysed event-related potentials (ERPs) time-locked to stimulus-onset and participants’ responses, focussing on attentional and motor-related components of the brain. In analysing the N1 component over electrodes P3/P4 and P5/P6, we first sought to understand whether attentional processes were affected by *Congruence, Turning Angle* and/or *Phase* ^72–74^. Due to the overlapping evoked potentials between the stimulus and response in our experiment, which we aimed to analyse separately, we utilised the Unfold Toolbox (Ehinger & Dimigen, 2019) for the deconvolution of all ERP components (see Methods).

The ANOVA of the mixed-effects model (R^2^ = 0.64) revealed significant main effects for the *Turning Angle* (F_2,187_ = 3.90; p = 0.0218;η*_p_*^2^ = 0.0400), but not for either *Congruence* (F_1,187_ = 0.10; p = 0.7429;η*_p_*^2^ = 0.0005), or the *Phase* (F_1,187_ = 0.26; p = 0.6099;η*_p_*^2^ = 0.0013). Post-hoc contrasts of *Turning Angles* (Figure 5D) only revealed significant differences for *Congruent* stimuli while *Turning*: 20° compared to 90° (p = 0.0449; d = 0.632), and between 45° and 90° (p = 0.0449; d = 0.638). For *Congruent* stimuli and *Straight* phase, we found no significance between 20° and 90° (p = 0.7564; d = 0.20), 20° and 45° (p = 0.7564; d = 0.04), nor between 45° and 90° (p = 0.4889; d = 0.25). This was also the case for *Incongruent* while *Turning* between 20° and 90° (p = 0.8767; d = 0.10), 20° and 45° (p = 0.8767; d = 0.12), and between 45° and 90° (p = 0.8767; d = 0.005)—and also for *Incongruent* and *Straight* phase between 20° and 90° (p = 0.5364; d = 0.12), 20° and 45° (p = 0.6051; d = 0.05), and between 45° and 90° (p = 0.6051; d = 0.07). These results indicate that the N1 amplitude decreases for increasingly sharper turns.

**Figure 5.**
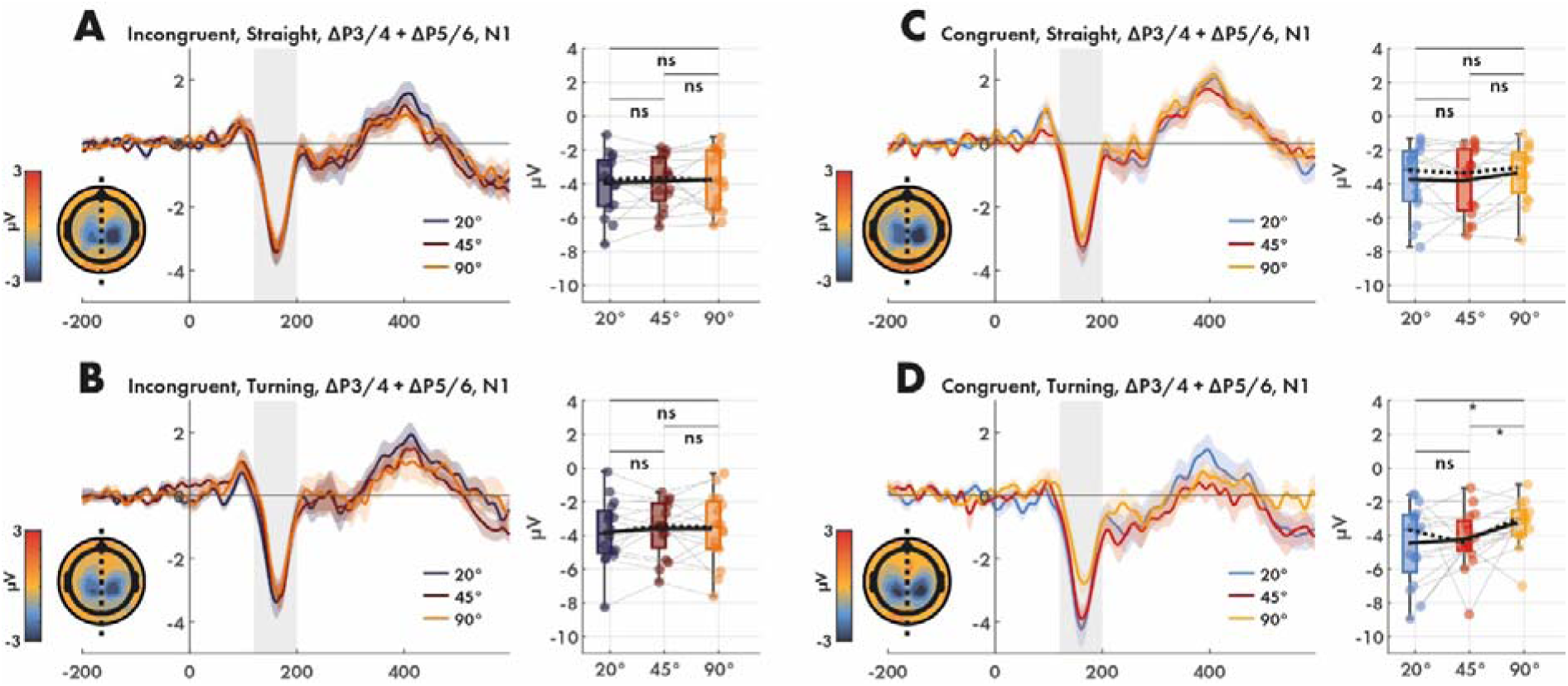
ERP plots of N1 over electrodes P3/4 and P5/6 between *Turning Angles* as grouped by *Congruence* and *Phases*. We only find differences for the *Congruent-Turning* between 20° and 90°, and between 45° and 90° corners. The amplitude within the grey areas were extracted and compared statistically in the adjacent raincloud plots. We used paired Wilcoxon Signed Rank tests with Benjamini-Hochberg correction for multiple comparisons. **A.** *Incongruent*-*Straight*. **B.** *Incongruent* -*Turning*. **C.** *Congruent*-*Straight*. **D.** *Congruent*-*Turning*.

We further analysed the P3 complex ^75^ over electrodes Pz, Cz and FCz to investigate whether the corners affected later attention-related potentials. The P3 complex is considered to reflect the allocation of attentional resources and cognitive processing associated with contextual features and context updating as well as target detection ^76^.

The mixed-effects ANOVA (R^2^ = 0.79) revealed main effects for *Turning Angles* (F_2,180_ = 7.87; p = 0.0005;η*_p_*^2^ = 0.0804), a tendency for *Congruence* (F_1,180_ = 2.82; p = 0.0943;η*_p_*^2^ = 0.0154), but not for *Phase* (F_1,180_ = 1.52; p = 0.2189; η*_p_*^2^ = 0.0083). We also identify an interaction effect among *Turning Angle* and *Congruence* (F_2,180_ = 4.79; p = 0.0093;η*_p_*^2^ = 0.0505) that required deeper investigation. Furthermore, we found a tendency between *Congruence* and *Phase* (F_1,180_ = 3.45; p = 0.0647;η*_p_*^2^ = 0.0188), but no interaction effects among *Turning Angle* and *Phase* (F_2,180_ = 1.07; p = 0.3444; η*_p_*^2^ = 0.0117) nor between all factors (F_2,180_ = 1.45; p = 0.2358;η*_p_*^2^ = 0.0159).

Post-hoc contrasts among *Turning Angles* (Figure 6A-D) revealed that only *Congruent* stimuli significantly differed. For *Congruent* stimuli displayed during the *Straight* phase, we found a significantly weaker P3 response for 90° as compared to 20° (p = 0.0057; d = 0.6811), and a tendency when compared to 45° (p = 0.0196; d = 0.5313). We also found that 45°-turns resulted in weaker P3 amplitudes compared to 20° (p = 0.0197; d = 0.5314). A similar pattern was found for *Congruent* stimuli during *Turning* phases: between 90° and 20° (p = 0.0012; d = 0.9251), 90° and 45° (p = 0.0437; d = 0.2323), and 45° and 20° (p = 0.0017; d = 0.7071).

**Figure 6.**
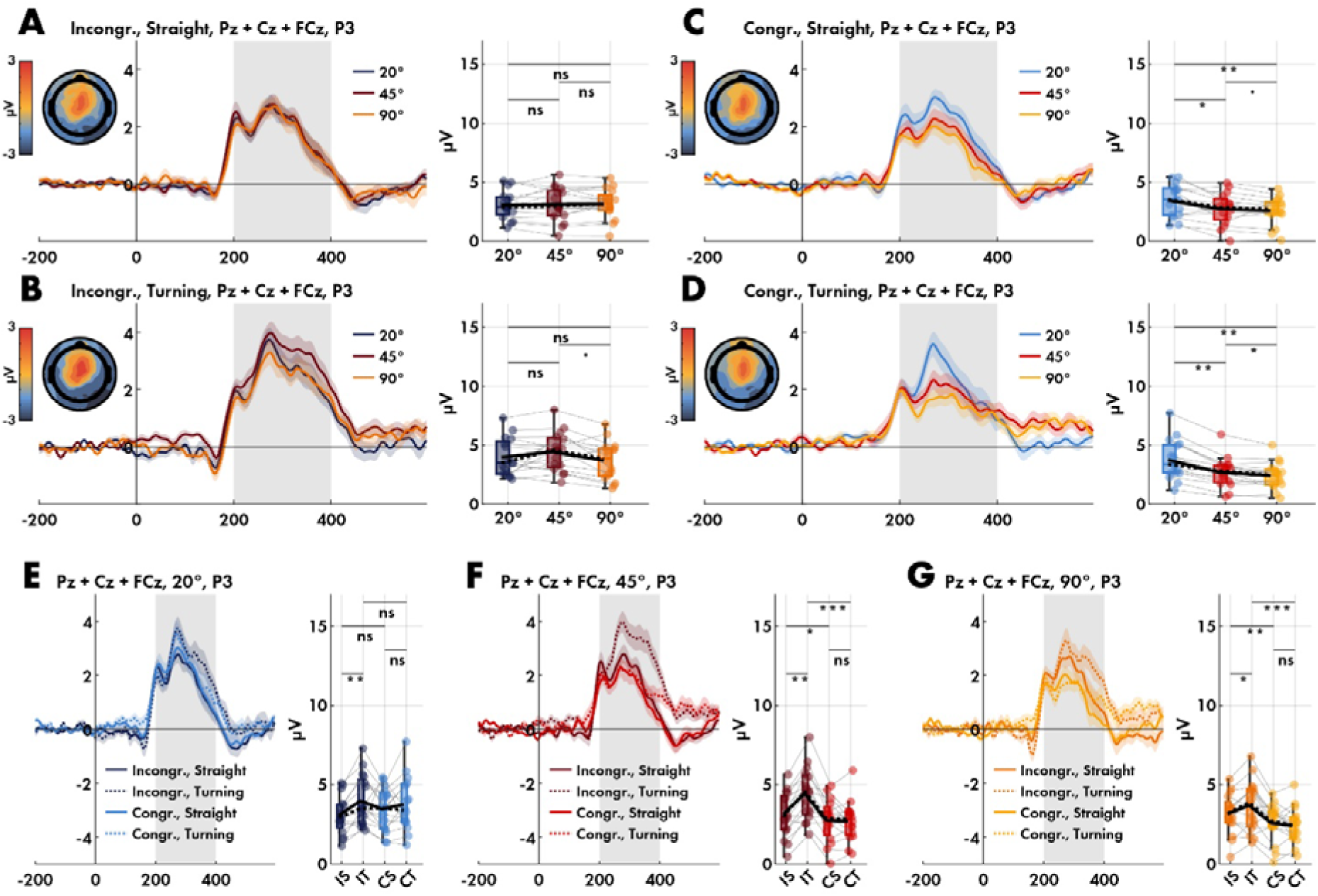
ERP plots of P3 over electrodes Pz, Cz, and FCz between *Turning Angles* as grouped by *Congruence* and *Phases*. We only find differences for the *Congruent* stimuli. The amplitude within the grey areas were extracted and compared statistically in the adjacent raincloud plots. We used paired Wilcoxon Signed Rank tests with Benjamini-Hochberg correction for multiple comparisons. **A.** *Incongruent* -*Straight*. **B.** *Incongruent* -*Turning*. **C.** *Congruent* -*Straight*. **D.** *Congruent* -*Turning*. **E-G.** ERP plots as grouped by the *Turning Angles* allowing for contrasting the *Congruence* with *Phases.* In a similar fashion, the raincloud plots are extractions of the positive peak amplitude in the grey area (200 and 400 ms). IS: *Incongruent* -*Straight Phase*. IT: *Incongruent* -*Turning*. CS: *Congruent* -*Straight*. CT: *Congruent*-*Turning*.

Due to the interaction effect, we also computed the remaining contrasts. For 20°-turns (Figure 6E), we only found a significantly stronger response for *Turning* compared to *Straight* for *Incongruent* stimuli (p = 0.0023; d = 0.6403). For 45°-turns (Figure 6F), we similarly found a significantly stronger response for *Turning* compared to *Straight* for *Incongruent* stimuli (p = 0.0038; d = 0.6569), but also between *Congruent* and *Incongruent* stimuli during *Straight* phases of the passages (p = 0.0084; d = 0.4855), and a strongly attenuated response during *Turning* compared to the *Straight* phase for *Congruent* stimuli (p < 0.0001; d = 0.8585). For passages with a 90° turn (Figure 6G), we again found a significantly stronger P3 response for the *Turning* as compared to the *Straight* phase for *Incongruent* stimuli (p = 0.0011; d = 0.8399). In addition, the contrast between *Congruent* and *Incongruent* stimuli during the *Straight* phase (p = 0.0229; d = 0.2820) and between *Incongruent* and *Congruent* during the *Turning* phase (p = 0.0004; d = 1.1852) both revealed significant differences. Finally, for the 90°-turn, we observe a similar pattern. A significant difference between *Straight* and *Turning* phases for *Incongruent* stimuli (p = 0.0386; d = 0.3637), a difference between *Congruent* and *Incongruent* stimuli during *Straight* phase (p = 0.0032; d = 0.4649), and a difference between *Incongruent* and *Congruent* stimuli during the *Turning* phase (p = 0.0004; d = 0.9195).

These results demonstrate that attentional processes are highly affected by the *Turning Angle*; we specifically found that the sharper the turn, the weaker the P3 responses. Also, *Congruence* and *Phase* affected the P3 responses to different extents. While more bodily torsion, i.e., *Turning*, resulted in stronger P3 responses, we observed that *Congruent* stimuli resulted in weaker responses.

## 3. Discussion

Although several scientists have voiced concerns about the limitations of static lab experiments, this remains the dominant approach in behavioural and cognitive neuroscience ^3–5,7,15,22,77^. The present study overcame the restrictions inherent in traditional brain imaging approaches by converting a static attention paradigm to a naturalistic version to investigate how active engagement with the environment affects the direction of attention and its impact on a behavioural and neuronal level. We used varying turning angles and directions of corners in a virtual built environment to implicitly direct spatial attention. Based on existing literature ^48,49,60,78^, we expected that directional changes during movement would lead to the allocation of attentional resources towards the direction of the turn based on the literature on saliency maps ^54^ and eye-tracking during turning behaviour ^79–81^, i.e. attention is directed towards the vanishing point of the anticipated path. We hypothesised (1) to find faster response time and higher accuracy for congruent trials, i.e., where stimuli are presented in the same visual hemifield as the turning directions, (2) to find slower response times and lower accuracy during the physical act of the turn compared to the in-and outlet of the passage, and (3) to find more pronounced early neural responses for *Congruent* stimuli, and more attenuated later neural responses for more physically demanding segments of the path.

Here we show that adapting a classic Posner paradigm to a naturalistic, movement-based protocol yielded strikingly different results. Contrary to our expectations, response times and accuracies were reversed in the naturalistic version of the attention cueing task, underscoring the critical importance of movement and ecological validity when studying human behaviour. Specifically, we found that during the physical act of turning, response times were significantly slower, and the number of incorrect responses increased compared to the straight phases before and after the turn. We also found that responses to stimuli displayed on the opposite side of the turning direction yielded significantly faster response times and higher accuracy. This flipped effect of the attentional cue indicates the involvement of early sensorimotor processes, possibly triggered by the active engagement with the built environment. Below, we provide an interpretation based on our behavioural and neuronal data.

The interaction with the environment naturally alters our visual field and motor processes ^62,63^. When considering only visual consequences, our findings are consistent with a driving study that also employed the Posner paradigm during turning behaviour. Their study also reported diminished detection performance during *Turning* phases compared to *Straight* phases and enhanced detection and faster response times for *Incongruent* stimuli compared to *Congruent* stimuli. While participants in our study engaged in full-body movement, compared to their movement-limited position during driving, their findings provide complementary support for our conclusions that movement through the environment and the associated change in optic flow lead to changes in attentional processes ^82^.

Our study included locomotion where existing research has shown that gaze behaviour relies on head-direction alignment with intended movement path, combined with eye-head coordination, to create a reference frame for controlling body motion through space ^83^. Furthermore, it has been shown that walking speeds decrease steadily as turning angles grow ^84–86^. It is also well established that while gaze direction typically leads head movement, it precedes the body’s rotational adjustments ^78^, which is likely responsible for the slowing of walking speed as observed in our data. This may be due to gaze and body segments adjusting earlier on more curved paths, and due to the fact that at turning points, gaze behaviour jumps ahead to preview distant sections of the path instead of following the usual sequence ^78^. In this sense, turning 90° corners affect us more and earlier as the predicted turn is greater.

Importantly, attention is directed towards the vanishing point of the expected path ^48,49^, which means the visual hemifields are impacted differently based on the direction of the turn. It suggests that turning to the left naturally shifts attention to the same side, and we note here that this side is precisely where novel contextual information needs to be processed. In contrast, the contralateral hemifield holds contextual information in the direction of the optic flow, which is therefore already known and thus predictable. Our interpretation is hence that when an observer turns a corner, causing a rotational optic flow ^87^, the visual field contralateral to the direction of rotation experiences an *outflow* of scene information, as objects move away from the centre of focus, creating a divergence of motion vectors. The hemifield in the turning direction, in contrast, experiences an *inflow* of information, where objects appear to approach the centre of rotation. This dynamic is essential for understanding how we perceive movement and navigate in turning environments ^82^. In our study, *Congruent* stimuli must be processed against a visual context that continuously expands and introduces novel contextual information (inflow of scene information) and thus itself requires effortful visual processing resulting in a higher cognitive load. The *Incongruent* stimuli, however, are presented against a visual context that is predictable as it is already known (outflow of scene information), resulting in smaller cognitive load. According to this view, one must expect to find better adaptation in *Incongruent* conditions as expressed by increased performance but also stronger early visual evoked responses. In contrast, more demanding phases of the corners, i.e., more acute-angled turns require more demanding motor execution leading to an augmentation of later cortical responses reflecting resource allocation ^71,88^.

Our behavioural data suggest that a stimulus presented within the predictable visual context, i.e. *Incongruent* stimuli, enhances action readiness, as indicated by faster response times and improved accuracy. Regarding response times, our model predicts a critical turning angle of 30°, where response times begin to be affected by the *Turning* behaviour. Although this value was derived from only three different turning angles in the present study, we emphasise that for tasks requiring attentional resources, it may be preferable to consider corners with angles less than 30°. This is also observed in our contrasts for *Congruence* (Figure 3F), where we only find significant differences for 45° and 90° turns. Consequently, we found that response times were only significantly affected by the turn if it reached 90° (Figure 3D), which aligns well with the existing literature as increasing motor demands results in depleted attentional resources and thus worse behavioural performance ^66,88–90^. In terms of accuracy, our model predicts a critical threshold at 15°, where *Congruence* begins to influence performance during a turn. Interestingly, we only find that performance varied for *Congruent* stimuli (Figure 4E), which according to our interpretation suggests that predictability of the visual context played a significant role. This is also suggested by the observed absence of difference during the *Straight* phases as compared to the *Turning* phase (Figure 4C). Our behavioural data suggest that stimuli appearing in the direction of ongoing movement impair attentional capacity as compared to the opposite site.

With regard to neuronal responses, existing research robustly showed that attended stimuli generate stronger N1 responses (for review; ^91^), suggesting that the N1 amplitude should be larger for stimuli on the same side as the direction of attention ^74,92–95^. In line with our interpretation on attention being impaired by the processing of novel information occurring on the same side as the turning direction, we observed weaker N1 responses for *Congruent* stimuli during the *Turning* behaviour. Accordingly, our data suggest that attention was reduced or insufficient for processing *Congruent* target stimuli embedded in the informative outflow patterns while *Turning* (Figure 5D)—however, we did not observe the same pattern for *Congruent* stimuli during the *Straight* phases of passages, which indicates that this effect may be tied to visual in-and outflow on the ipsi-and contralateral turning sides when moving through the turn. This may be due to the motoric role of the pulvinar and superior colliculus in orienting behaviour ^96,97^, as well as the saliency maps in the superior colliculus ^98–100^, which has strong connections with the pulvinar ^27,101^. Their connections with the thalamic reticular nucleus allow for selective gating of visual information during movement ^102,103^. Reduced N1 amplitudes during *Turning* could reflect this subcortical filtering of visual inputs to prioritise motor control as reflected in impaired attentional performance.

This is further reflected in the late P3 complex, which originates from stimulus-driven attention mechanisms and reflects belief updating and integration of environmental changes ^75,76,104^. Our data suggests that P3 systematically decreases in amplitude with increasing *Turning Angles* for *Congruent* stimuli, but not for *Incongruent* stimuli, which aligns with our behavioural results (Figure 6A-D). This supports the concurrent interpretation of attentional processes being affected by the motor-related demands during physical turning. The 90° corner necessitates more complex motor planning and execution, leading to larger sensorimotor demands, which in turn is reflected in reduced P3 amplitudes. This aligns with recent findings demonstrating decreased P3 amplitudes over central areas with increasing motor demands ^71,105^. Post-hoc contrasts between *Phase* and *Congruence* revealed large effects between *Incongruent* and *Congruent* stimuli regardless of *Phase* for both 45° and 90°. However, we only observed significant effects between *Turning* and *Straight* phases of passages for *Incongruent* stimuli, but not for *Congruent* stimuli (Figure 6E-G). These results suggest that the phase is irrelevant for turns under 20°; beyond this, the difference between *Straight* and *Turning* segments becomes significant due to the congruency effect during the turn. Thus, a 20° turn seems similar to a *Straight* segment. Altogether, these results support the idea that P3 amplitude reductions are linked to increased cognitive load when motor tasks compete for attentional resources ^71,88^. Though the present study did not record neural data from these deeper sources, a likely explanation for the observed patterns is that increased task difficulty alters the information flow in pulvino-cortical connections, affecting the underlying mechanisms responsible for P3 generation.

These findings raise profound implications regarding the reciprocal relationship between built environments and human cognition, while simultaneously highlighting the critical importance of naturalistic paradigms in cognitive neuroscience. From an architectural perspective, the fact that numerous cities are designed based on grids that exclusively present 90° turns ^9^ might have larger cognitive load than expected on city-dwellers ^106^—and even more so for the less agile inhabitants, such as the elderly ^107^, people with Alzheimer’s ^108^, and people with Parkinson’s ^109^. These results suggest a need to reconsider city structures. From a scientific perspective, the apparent disparities between static and naturalistic conditions suggest we should be cautious about generalising conclusions from constrained experiments into the dynamic, embodied nature of behaviour and cognition in the real world. The unconscious influence of architectural features on attention and behaviour suggests that our constructed spaces may have inadvertently contributed to the shaping of cognition throughout human history, yet our understanding has been limited by static experimental paradigms. Given that the brain’s neural mechanisms did not evolve in static settings ^3,22,110–112^, but rather adapted to handle dynamic scenarios like approach/avoidance decisions and obstacle navigation, empirical investigation through large-scale naturalistic experimentation is essential. Our cognitive abilities are fundamentally built upon sensorimotor strategies inherent in such dynamic behaviours ^113^, making the study of cognition in real-world contexts crucial for understanding how both evolutionary history and built environments shape human behaviour and cognitive processing.

Our study brings evidence that human behaviour is more susceptible to environmental features and contextual information than static experimentation anticipates, underlining the importance of naturalistic cognitive neuroscientific experiments ^14,16^. Attention research has a long tradition in behavioural and neuroscience research and we thus adapted a robust attention paradigm, i.e., Posner paradigm ^17^, by using corners in the built environment to pre-orient the body and the attentional capacities. To our surprise, our study yielded a flipped effect on attention, leading us to revise our initial hypothesis. While we found that participants’ response times and accuracy significantly worsened during the physical act of turning a corner, we also observed that stimuli presented on the same side as the turning direction led to notably longer response times and a higher number of incorrect responses. Neural responses suggest that both the N1 and P3 components were affected by both visual and motor processes, particularly during more challenging conditions like 90° turns. While previous research has shown that N1 can index attention by demonstrating its sensitivity to visual context under static conditions, our findings extend this by showing how built environmental context critically modulates these attentional effects. Similarly, for the P3, we suspect that the organisational structure of the pulvinar and its functional relationship with cortical areas have influenced attention in favour of stimuli appearing in the hemifield contralateral to the turning direction. This is supported by another study on turning behaviour reporting that walking along a curved path is associated with a contralateral shift of activation in the basal ganglia ^114^.

We take these results as an appeal to a paradigm-shift in cognitive neuroscience in favour of naturalistic paradigms, challenging the long-standing dominance of static paradigms in cognitive neuroscience. If the embodied brain responds so dramatically differently in mobile situations, what other fundamental assumptions about cognition might be overturned when we step out of the lab?

## 4. Methods

### a. Participants

Twenty participants (9 females), aged between 23 and 54 years (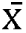= 30.1, σ = 7.18), consented and were recruited for the experiment that took place in Berlin Mobile Brain/Body Imaging Lab (BeMoBIL) at Technical University of Berlin. The experiment was authorised by the local ethics committee. In the 120 sqm VR space, they completed a naturalistic attention task in which the built environment and body posture acted as implicit cues. During corner turns, 10 small spheres appeared either on the same side (*Congruent*) or the opposite side of the turn (*Incongruent*). These spheres were triggered by spatially fixed, invisible anchors, making their appearance dependent on location rather than time ^115^ (Figure 7). Each participant completed 240 trials, with each trial consisting of 10 pseudo-randomly presented stimuli (5 ipsilateral and 5 contralateral). All instructions were presented in the VR space, minimising the interaction between researchers and participants. Participants correctly identified 90.38% of the stimuli, with an error rate of 9.62%. After data preprocessing, a total of 37,882 valid trials remained for analysis. We refer to stimuli ipsi-and contralateral to the turn as *Congruence* (2 levels: *Congruent* and *Incongruent*, respectively); 20°, 45°, and 90° corners as *Conditions* (3 levels); stimuli from 1 to 3 and 8 to 10 were classified as *Straight* and stimuli from 4 to 7 as *Turn*. Both *Straight* and *Turn* are referred to as *Phases* (2 levels).

**Figure 7.**
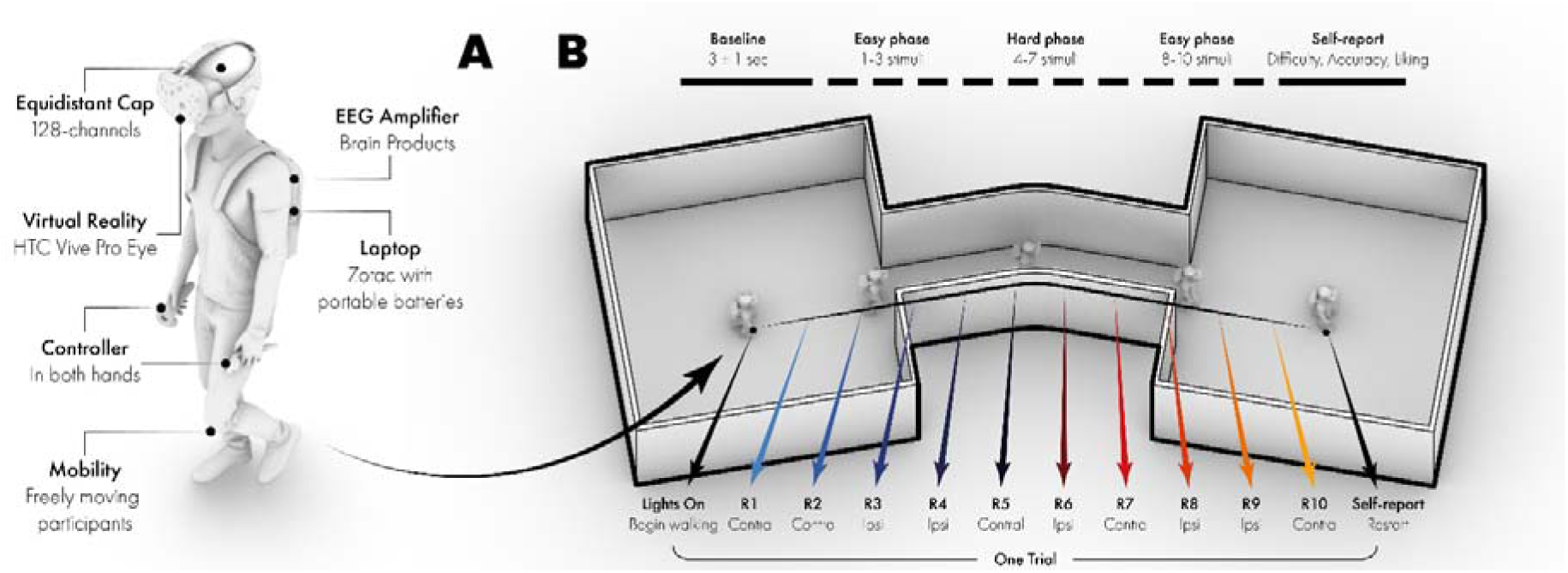
**A.** The equipment setup for each freely moving participant. **B.** The structure of a single trial. The dashed line above illustrates the how the baseline preceded the walking, and how the *Phases* and *Congruence* were categorised. The isometric perspective of the corner is an example of 20° corner towards the right. The responses, named below as R1 through R10 refer to responses, alternates pseudo-randomly between ipsilateral and contralateral stimuli.

### b. Equipment

For Virtual Reality (VR), we used the HTC Vive Pro Eye headset, equipped with dual OLED displays providing a combined resolution of 2880 x 1600 pixels (1440 x 1600 per eye) at a refresh rate of 90 Hz and a field of view of approximately 110 degrees. The VR experiment was programmed on Unity (version 2019.2) and ran on a Zotac (MAGNUS EN1060), equipped with a GTX 1060 (NVIDIA GeForce), and custom-made batteries to make the setup entirely mobile. Electroencephalography (EEG) data were collected from 128 active wet electrodes at a sampling rate of 1,000 Hz and a high-pass filter of 0.016 Hz and a low-pass filter of 250 Hz (BrainAmp Move System, Brain Products, Gilching, Germany) using an elastic cap with an equidistant layout (EASYCAP, Herrsching, Germany). All reported electrodes are based on the electrode nearest to the standard locations of the 5% electrode system for EEG. The reference electrode was located near F3 while the ground electrode was located near F4 and impedance for all electrodes was brought below 15 kΩ. All data was streamed and recorded using Lab Streaming Layer ^116^.

### c. Experimental paradigm

After equipping participants with the mobile EEG amplifier, 128-channel cap, VR goggles, and handing them a controller in each hand, they all underwent a training phase in VR to ensure they understood their task. The VR spaces were designed based Space Syntax analyses (Figure 1A and 1B), which simulates how spatial configurations of buildings and urban environments presents themselves along a path ^9,117,118^. A full trial consists of starting in a pitch darkness for 3 seconds (±1 second). Lights will turn on and the participants will be facing a corner that either bends 20°, 45°, or 90° to either the right or left side. They are instructed to start passing through the corridor as soon as the lights turn on. While they are walking, small orange spheres are popping up on either the right or left visual field (20° off the centre). Once the stimuli appear, they are instructed to click the corresponding controller as fast as they can, i.e. stimuli appearing on the right resulted in clicking the right controller. Once they get to the end, they are asked to assess the trial on difficulty and accuracy. They then turn around to face the corridor again and click the controller to move along to the next trial (Figure 7). Additionally, we note that real-world scenarios are not necessarily tailored to specific tasks, we thus focus on using turning direction as cues that appear with 50/50 chance of indicating target location, which creates purely exogenous attentional capture ^17,119,120^.

### d. Behavioural data

We extracted both response times and correct responses as measures of the behavioural performance for the different turns. Response times were computed as the difference between stimulus-onset and response-event, while the correct responses were collected by matching the condition and the responding controller. The modelling of the behavioural data was based on linear mixed-effects models, which were then submitted to an ANOVA. All post-hoc comparisons were corrected for multiple comparisons where appropriate, using Benjamini-Hochberg (alpha: 0.05). We only used paired t-tests when the data did not violate Levene’s test nor Lilliefors normality test. In such cases, we used Wilcoxon Signed Rank test. All data was processed in MATLAB v. R2024b ^121^.

### e. EEG Analysis

#### i. Preprocessing

For analysing our data, we used EEGLAB (v. 2023.1; ^122^), which is a plugin to MATLAB v. R2024b ^121^. We applied the BeMoBIL Pipeline ^123^, which consists of down-sampling the data to 250 Hz, removing noisy parts of the dataset, interpolating channels that are deemed bad based on spectra, kurtosis and probability features (threshold = 4), and finally filtering each dataset twice; one version is bandpass filtered between 2.5 Hz and 100 Hz ^124^, and another is bandpass filtered between 0.2 Hz and 40 Hz, which is more suitable for ERP analyses. Channels were referenced to the average reference. The dataset with the broader bandpass filter underwent Adaptive Mixture Independent Component Analysis (AMICA; ^125^), where the resultant ICA spheres and weight matrices were applied to the dataset with the narrower bandpass filter. Independent components associated with eye movements, such as blinks and horizontal movements, were classified using ICLabel ^v. 1.4; 126^ and subsequently removed.

As the nature of the experiment guarantees overlapping evoked potentials between the stimulus and the response, which we wanted to analyse separately, we applied the Unfold Toolbox ^127^. The toolbox is designed to analyse overlapping ERPs in EEG data, using general linear models (GLMs) to deconvolve and estimate contributions from multiple rapidly occurring events. It models non-linear effects and interactions between factors through time-expanded techniques, i.e. time-shifted design matrix on which a GLM will be fit for each time-point, resulting in a linear deconvolution. ERPs were epoched-200 ms before and 600 ms after the stimulus onset. Unfold’s automatic artefact rejection resulted in removing 20.02% of the data, corresponding to a total of 42,641 trials (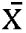= 2244 trials/participant, σ = 398 trials). We then modelled the data using two formulas to separate the stimulus response from the motor response. Using the beta for each timepoint, we retrieved the separated ERPs based on following formulas:

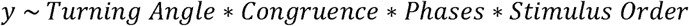

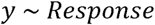

This attempts to model the ERPs based on: *Turning Angles* (20°, 45°, 90°), *Congruence* (Congruent, Incongruent), *Phase* (Turning, Straight), and *Stimulus Position* (1 through 10).

#### ii. Event-related potentials

For all ERPs, baseline was set to 200 ms before stimulus onset to stimulus onset. For the N1, we recovered the ERP using the estimated beta values from Unfold. Then we subtracted the electrodes ipsilateral and contralateral to the stimulus direction. Finally, we automatically extracted the negative peaks, using a simple peak-detector, in the time window between 120 and 200 ms over channels P3/4 and P5/6 and then averaged between the electrodes. For the P3, we also recovered the ERP using Unfold and automatically extracted the positive peaks between 200 and 400 ms over channels Pz, Cz, and FCz and then averaged between the electrodes. All post-hoc comparisons were computing the Wilcoxon Signed Rank Test, and all multiple comparisons have been corrected using Benjamini Hochberg (alpha: 0.05). For all mixed-effects regressions, we computed both with and without interactions and selected the approach that best fitted the data based on AIC, BIC, and log-likelihood.

